# Secreted HLA Fc-Fusion Profiles Immunopeptidome in Hypoxic PDAC and Cellular Senescence

**DOI:** 10.1101/2023.04.10.536290

**Authors:** Nicholas J. Rettko, Lisa L. Kirkemo, James A. Wells

## Abstract

Human leukocyte antigens (HLA) display peptides largely from intracellular proteins on the surface of cells in major histocompatibility complex (MHC)-peptide complexes. These complexes provide a biological window into the cell, and peptides derived from disease-associated antigens can serve as biomarkers and therapeutic targets. Thus, proper identification of peptides and the corresponding presenting HLA allele in disease phenotypes is important for the design and execution of therapeutic strategies using engineered T-cell receptors or antibodies. Yet, current mass spectrometry methods for profiling the immunopeptidome typically require large and complex sample inputs, complicating the study of several disease phenotypes and lowering the confidence of both peptide and allele identification. Here, we describe a novel secreted HLA (sHLA) Fc-fusion construct that allows for simple peptide identification from single HLA alleles in two important disease models: hypoxic pancreatic ductal adenocarcinoma (PDAC) and cellular senescence. We identify hypoxia and senescence-associated peptides that could act as future targets for immunotherapy. More generally, the method streamlines the time between sample preparation and injection from days to hours, yielding allele-restricted target identification in a temporally controlled manner. Overall, this method identified >30,000 unique HLA-associated peptides across two different HLA alleles and seven cell lines. Notably, ∼9,300 of these unique HLA-associated peptides had previously not been identified in the Immune Epitope Database. We believe the sHLA Fc-fusion capture technology will accelerate the study of the immunopeptidome as therapeutic interest in HLA-peptide complexes increases in cancer and beyond.

## INTRODUCTION

Human leukocyte antigen (HLA) molecules display peptides derived from largely intracellular proteins on the surface of somatic cells in major histocompatibility complex (MHC)-peptide complexes^1^. The pool of HLA-associated peptides, collectively known as the immunopeptidome, can reflect the proteome of a cell in a given phenotype^2,3^. Consequently, peptides derived from mutated oncoproteins or disease-associated antigens in MHC-peptide complexes can serve as phenotypic biomarkers and targets for immunotherapies. Over the past decade, antibodies targeting MHC-peptide complexes containing peptides from tumor-associated antigens, such as Wilms tumor protein (WT1) and p53^R175H^, have demonstrated cytotoxic potential^4,5^

Given the therapeutic promise of targeting MHC-peptide complexes, profiling the immunopeptidome in disease phenotypes is critical for identification of biomarkers and the downstream development of therapeutics. To date, the most common way to broadly identify these peptides is through LC-MS/MS analysis. Yet, challenges remain with current immunopeptidomic methods, including large sample input (up to a billion cells)^6^, contamination^7^, cell adherence^8^, and importantly, immunoprecipitation of all class I MHC-peptide complexes regardless of allele—up to six per cell^9^. As HLA molecules are the most polymorphic genes – with upwards of 10,000 different alleles in the human population – and with each allele accommodating different peptide motifs, the common procedure of pan-MHC immunoprecipitation can yield tremendously complex peptide mixtures. This complexity is further amplified as many peptides can bind multiple HLA molecules^10^, making it difficult to deconvolute which HLA molecule is presenting which peptide—a critical basis for antigen design and therapeutic development. Additionally, as HLA-associated peptides are inherently difficult to analyze due to similarity in size and amino acid composition^7^, multi-allelic peptide samples can compound complexity and decrease confidence in peptide origin and sequence.

To circumvent these challenges, some have utilized engineered cell lines to express single HLA constructs. Generation of membrane bound mono-allelic HLA cell lines, such as B721.221, have had success in identifying tens of thousands of HLA-associated peptides^10,11^. While this method produces peptide samples displayed by a specific HLA, this approach is limited to cell lines that are inherently HLA-null, thereby restricting the breadth of cell types and biological contexts that can be surveyed. Additionally, these cell lines have been reported to have low, but residual, endogenous HLA expression, potentially contaminating the samples^7^. Others have developed soluble HLA constructs lacking the transmembrane domain, allowing the HLA to be loaded and subsequently secreted into the milieu to be purified through affinity tags^12^. While this method retains allele specificity and can be expanded to different cell types, it remains unoptimized and typically relies on large bioreactors for cell culture^11,12^. Additionally, these soluble HLA methods still require copious amounts of material (10-25 mg) as it demands several downstream purification and fractionation steps^12,13^. Due to these various factors, current mono-allelic approaches are not amenable for identifying disease-specific immunopeptidomes.

Here, we describe a streamlined mass spectrometry-based method based on a doxycycline-induced secreted HLA Fc-fusion construct (sHLA-Fc fusion). We apply the sHLA-Fc fusion capture technology to profile the immunopeptidome of two important cellular states, hypoxia and senescence. We show the method provides pure, temporally-controlled, mono-allelic samples without requiring cell lysis or peptide fractionation, and can be prepared in a matter of hours as opposed to the 2-3 days required by typical immonopeptidomic methods^7^. Using samples ranging from 12-130 million cells, we identified >30,000 peptides across 7 cell lines and 2 HLA alleles ranging from ∼600 to 10,000 peptides per sample depending on cell line and condition. We identified unique phenotype-restricted HLA-associated peptides in both hypoxic and senescent cells. We believe the sHLA Fc-fusion capture technology will accelerate profiling the immunopeptidome in disease states, which will be paramount towards continued novel target discovery and therapeutic development.

## RESULTS

Fc-fusion proteins have served as structural scaffolds for increased expression, solubilization, and purification of the extracellular domain of membrane proteins^14,15^. As HLA proteins lacking the transmembrane domain have been shown to be properly folded and loaded with peptide cargo^12,13^, we hypothesized that a HLA Fc-fusion (henceforth referred to as sHLA) could be loaded and secreted similarly (**Figure 1A**). With mono-allelic B721.221 cell line datasets providing a standard dataset for peptide processing and binding prediction^10,11^, we sought to determine if our sHLA Fc-fusion could capture HLA-associated peptides in this established cell model. We engineered two B721.221 cells, each transduced to express a sHLA Fc-fusion of either HLA-A*02:01 and HLA-B*35:01, two of the most common alleles present in 39% and 8% of global population, respectively (www.iedb.org), under doxycycline induction. After 52 hours with or without doxycycline treatment, media was collected and sHLA proteins were immunoprecipitated using magnetic Protein A beads and analyzed by western blot (**Figure 1A&C**). As western blotting indicated successful secretion of the sHLA Fc-fusion complex from B721.221 cells, we subsequently isolated the sHLA-containing media from doxycycline treated B721.221 cells and performed LC-MS/MS on the peptides derived from the sHLA Fc-fusion complexes through acid elution and standard desalting procedures. Data were analyzed using a stringent 1% FDR, and m/z spectra demonstrated clean +1 and +2 charge populations (**Figure 1B**). The peptide search (PEAKS Online) identified consensus peptides in close agreement with peptide lengths (**Figures 1E&F**), dominant anchor residues (**Figures 1G&H**), and affinity prediction profiles (**Figures 1I&J**) of previously reported membrane bound HLA for each allele^11^.

**Figure 1:**
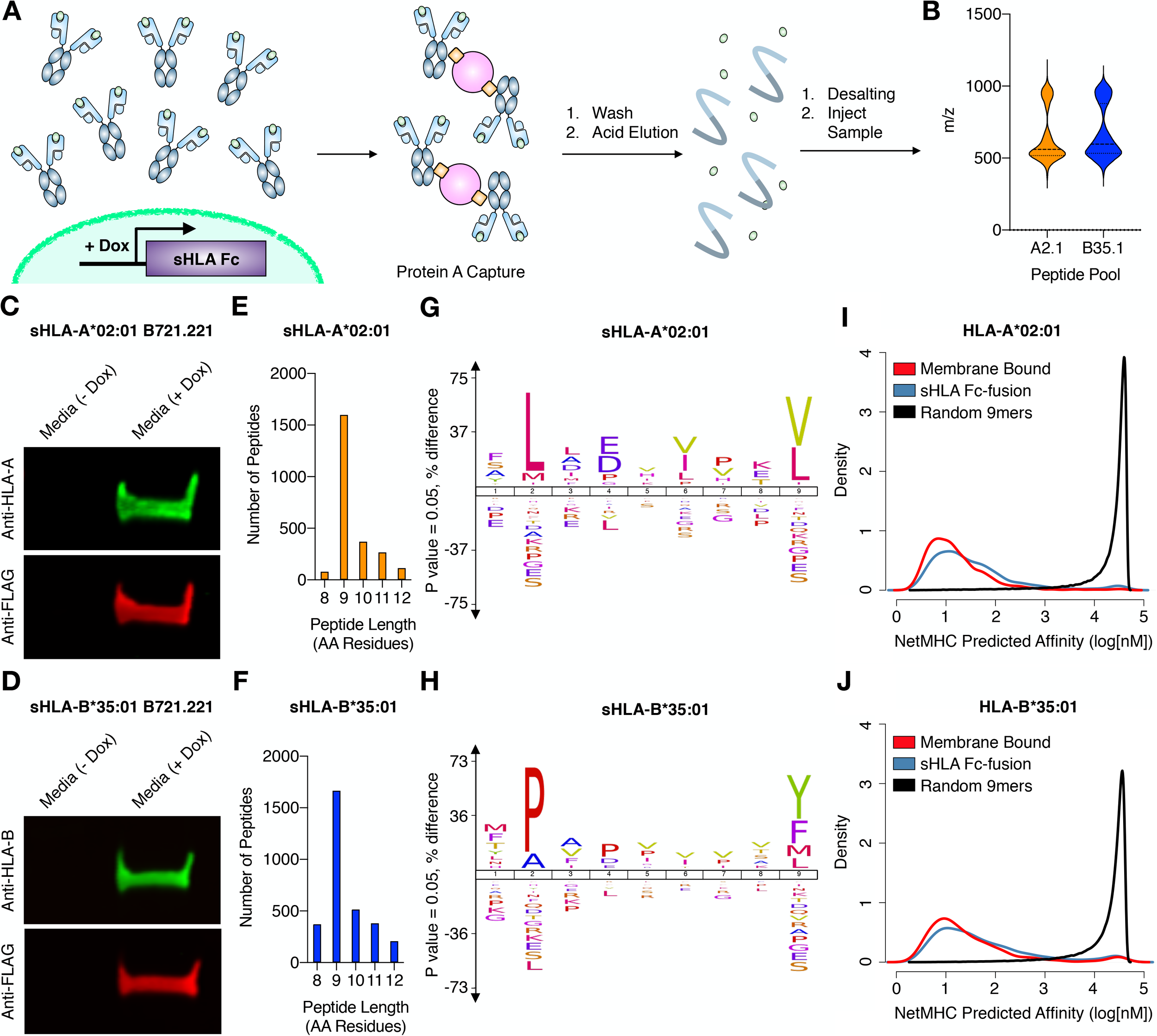
Workflow for sHLA cell line generation and subsequent immunopeptidomics. (A) Secreted HLA (sHLA) Fc-fusion is virally transduced into cells of interest. sHLA is induced using doxycycline after cells have undergone disease-specific perturbation. The media from cells is isolated and sHLA are immunoprecipitated, washed, and the peptides are eluted for analysis by LC-MS/MS. (B) Mass:charge of identified peptides from sHLA mono allelic B721.221 cell lines (n=2). (C &D) Western blot of eluted sHLA Fc-fusion protein captured from doxycycline-treated or –free media of sHLA monoallelic B721.221 cell lines expressing sHLA-A*02:01 or sHLA-B*35:01, respectively. (E & F) Quantification of peptide length from cell lines expressing sHLA-A*02:01 or sHLA-B*35:01, respectively. (G & H) ICE logos of 9mer peptides from cell lines expressing sHLA-A*02:01 or sHLA-B*35:01, respectively. (I & J) NetMHC predicted affinities of the 9mer peptides from sHLA-A*02:01 or sHLA-B*35:01 immunopeptidomics datasets, respectively, compared to published 9mers identified from membrane bound monoallelic B721.221 cells and a published list of 100,000 9mer peptides^10,11^.

Hypoxia and cellular senescence are two disease-phenotypes in which biomarkers are of interest, yet little is known regarding the respective immunopeptidome. We chose three cell lines commonly used in studying these phenotypes and transduced them with either our sHLA-A*02:01 or sHLA-B*35:01 expression constructs. For profiling hypoxia in PDAC, cells were grown for 5 days either under 20% O_2_ (normoxia) or 1% O_2_ (hypoxia), with doxycycline-treatment beginning after 3 days (**Figure 2A**). Establishment of the hypoxic phenotype was confirmed by Glut1 expression (**Figure 2B**). For profiling cellular senescence, cells were treated with either DMSO or 250 nM doxorubicin for 24 hours to generate growing and senescent samples. Doxycycline-treatment immediately followed DMSO treatment in growing samples and 8 days post-doxorubicin treatment in senescent samples (**Figure 2A**). The senescence phenotype was confirmed by beta-galactosidase activity staining (**Figure 2C**). Each conditioned cell line was prepared and assayed across four biological replicates, increasing the confidence in the presented peptide identifications.

**Figure 2:**
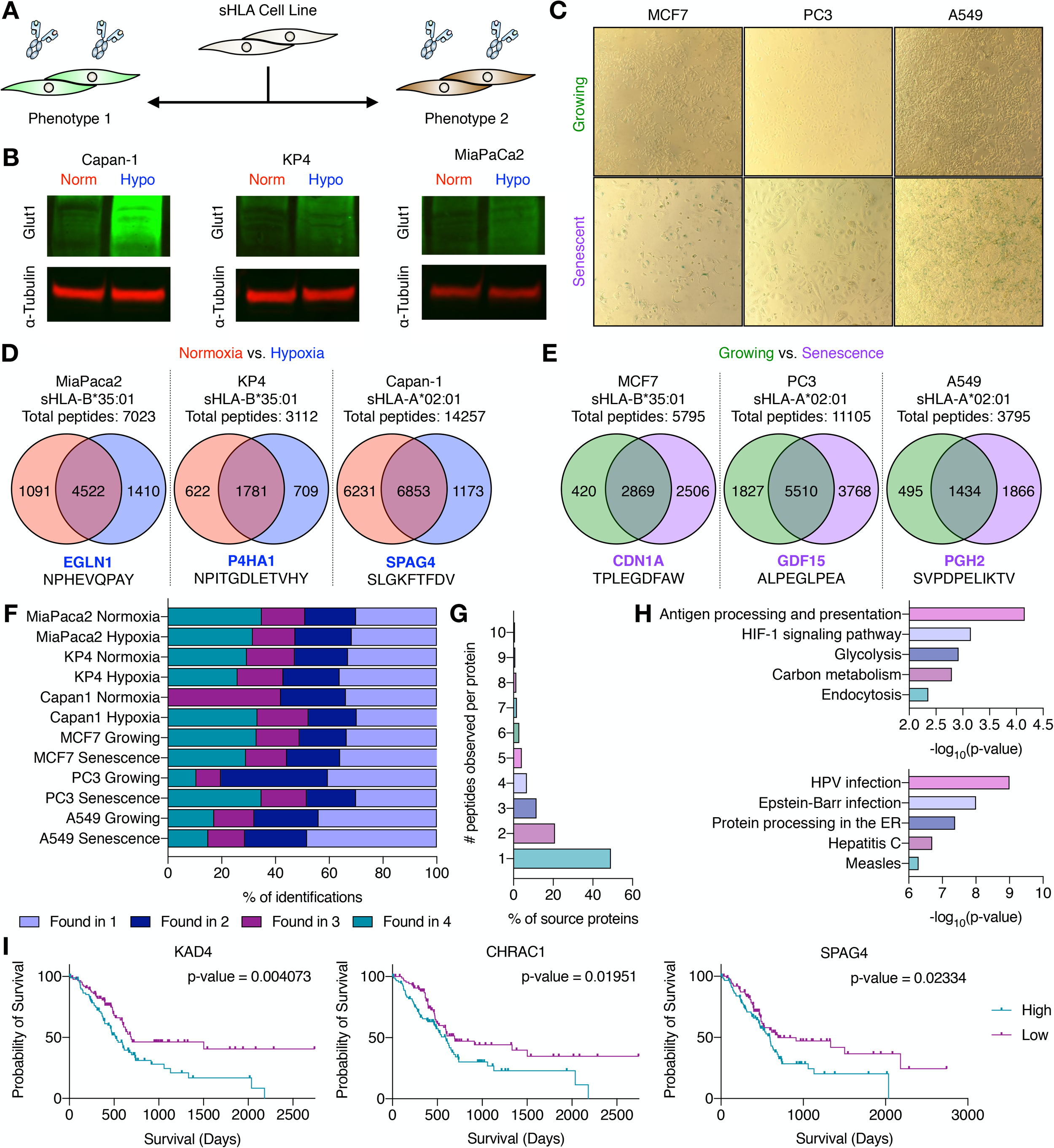
Immunopeptidomics of hypoxic and senescent sHLA cell lines. (A) Strategy for identifying phenotype-specific HLA-associated peptides. (B) Western blot for GLUT1 expression in sHLA cell line samples under normoxic and hypoxic conditions. (C) β-galactosidase staining of sHLA cell line samples. (D & E) Venn diagrams of identified peptides across sHLA cell lines across all replicates (n=4, n=3 for CAPAN-1 Normoxia) with a highlighted high-confidence peptide found to be phenotype-specific from each cell line. (F) Reproducibility of peptide identification for each sHLA cell line under each condition across all replicates. (G) Number of peptides identified per protein in CAPAN-1 normoxic samples. (H) GO-term enrichment of select biological processes in high-confidence peptides for hypoxia (top) and senescence (bottom) datasets. (I) Survival curves for proteins of which high-confidence hypoxia-specific peptides were identified from sHLA CAPAN-1 cell lines.

Across normoxic and hypoxic samples, we identified 7,023 unique peptides in MiaPaCa-2, 3,112 unique peptides in KP4, and 14,257 unique peptides in Capan-1 cells. Peptide identifications ranged from ∼600-10,000 per sample across the different phenotypes, with Capan-1 normoxic samples yielding nearly 10,000 HLA-A*02:01-associated peptides from a ∼130 million cell sample (**Figure 2D**). This proteomic depth demonstrates the broad applicability of the method towards profiling the immunopeptidome of a single allele. We identified 99, 61, and 135 peptides unique to hypoxia with high confidence in at least 3 out of 4 hypoxic biological replicate samples in MiaPaCa-2, KP4, and Capan-1 cells, respectively. A number of these hypoxia-specific peptides, such as EGLN1 (HLA-B*35:01: NPHEVQPAY) or P4HA1 (HLA-B*35:01: NPITGHLEHTVHY) are known to interact with or be driven by the canonical hypoxia-inducible transcription factors (HIF, **Figure 2D**)^16,17^. Excitingly, from these high confidence hypoxia-associated peptides, we identified peptides from several biomarkers of hypoxia, including KAD4 (HLA-A*02:01: NLDFNPPHV)^18^, DYH11 (HLA-A*02:01: VLFEDAMQHV)^19^, and SPAG4 (HLA-A*02:01: SLGKFTFDV)^20^. Interestingly, there were peptides found with high confidence exclusively in hypoxic samples, such as the HLA-A*02:01-associated KLHGGILRI derived from EGLN3, despite other peptides from the same protein being identified in normoxic samples. While these data suggest some differential processing in hypoxia versus normoxia, the lack of signal in the normoxic condition does not preclude it being present in the normoxic context at a level not detected. Sensitivity of the mass spectrometer or signal interference from other peptides could both play a role in explaining absence of signal in one context versus another. More work will be required to dissect the underlying pathways that are over-represented in the immunopeptidome of hypoxia and could serve as future therapeutic targets for treating solid tumors.

Across growing and senescent samples, we identified 5,795 unique peptides in MCF7’s, 11,105 unique peptides in PC3’s, and 3,795 unique peptides in A549 cells (**Figure 2E**). Of the peptides identified exclusively in senescence within each cell line, 466 in MCF7, 1021 in PC3, 190 in A549, were found in at least 3 out of 4 biological replicates, providing confidence in these identifications. Excitingly, from these high confidence peptides exclusive to senescence, we identified peptides from senescence-associated secretory phenotype (SASP) biomarkers, including GDF15 (HLA-A*02:01: ALPEGLPEA) and PGH2 (HLA-A*02:01: SVPDPELIKTV)^21,22^. Additionally, a peptide derived from the senescence marker p21 (CDN1A)^23^, was found exclusively in senescent MCF7 samples with high confidence, further showcasing the ability of the method for isolating phenotype-relevant peptides (**Figure 2E**). Remarkably, these large datasets for MCF7 senescent samples were collected using less than 17 million cells per sample with an average peptide ID count of ∼3,200. This highlights the potential of using this method with smaller cell samples without sacrificing peptide counts. Interestingly, we saw a reproducible increase in peptide identifications across senescent samples compared to their growing counterparts. As senescent cells have an active secretory phenotype, our sHLA construct may be advantageous and have improved trafficking in cells and phenotypes with increased secretory activity.

Evaluating the robustness in immunopeptidomic experiments is paramount to building high-value datasets. Our sHLA technology showcases high levels of reproducibility, with the majority of peptides being identified in at least 2 biological replicates per condition (**Figure 2F**). A majority of the peptides identified were the sole peptide derivative from a given protein sequence, further highlighting the need for highly reproducible datasets to increase confidence **(Figure 2G**). When focusing on high confidence peptides found exclusively in either the hypoxic or senescent datasets, we identified biological processes that recapitulate the biological perturbation (**Figure 2H**). Specifically, the high confidence peptides identified in the hypoxic datasets enrich for biological processes indicative of HIF-1 activation, such as glycolysis and carbon metabolism – all of which are critical in response to hypoxic stress. Likewise, high confidence peptides in the senescent datasets yielded pathways associated with infection and inflammation – biological processes intimately linked with senescence^24^. Finally, we show that not only do our high confidence peptides enrich for disease-associated biological processes, they may also be indicators of disease prognosis that could serve as useful biomarkers in detection and/or therapeutic intervention with further validation **(Figure 2I**).

Overall, across all cell lines our sHLA Fc-fusion immunopeptidomic method identified more than 30,000 peptides. Of these peptides, when cross-referenced with the Immune Epitope Database (www.iedb.com), >9,000 were new to the database. Hence, this sHLA technology profiles the immunopeptidome of a given allele with immense depth that can be useful in candidate identification as well as informing predictive algorithms for peptide binding and generation.

## DISCUSSION

Our results show that the sHLA Fc-fusion technology has some significant advantages for immunopeptidomics. Using allele-defined sHLA Fc-fusions, it is possible to generate reproducible immunopeptidomes that align with consensus sequences found for membrane bound forms using more laborious ubiquitous pull-downs or cells engineered to express only one specific HLA. Robust samples can be produced from fewer cells and can be processed in hours as opposed to days. Doxycycline-induction allows one to capture discrete time windows and particular cellular states as shown here for hypoxia and senescence. These advantages allow one to profile disease models including, but not limited to, viral or bacterial infection, chemical perturbation, or oncogenic transformation. High order and complex models, such as co-culturing different cells and profiling the immunopeptidome of a single allele in only the cell line of interest, are also possible applications of this sHLA technology that may be hindered with commonly used approaches. Lastly, sHLA datasets can inform peptide binding and generation prediction algorithms that will be important as more HLA alleles are characterized.

While the sHLA technology shows advantages, there are limitations. Since it requires lentiviral transduction to generate stable cell lines, primary cell lines or those excised directly from patients may not be amenable to lentiviral perturbation or *ex vivo* passaging. Although the consensus motifs for sHLAs qualitatively match those published from membrane bound HLAs, due to conditional variability in sHLA expression more extensive platform development would be required for absolute or relative quantification. We have also seen evidence that the sHLA Fc-fusion may compete with the endogenous membrane bound HLA for peptide loading, suppressing folding and secretion. For example, HLA-A*02:01 sHLA constructs were suppressed in expression in HLA-A*02:01^+^ IMR90 fibroblasts and Panc-1 cells (data not shown). In these cases it may be possible to knock out the endogenous allele to avoid competitive loading effects between membrane bound and the exogenous sHLA construct.

From the proteomic depth we attained, it was possible to capture numerous peptides per protein. Interestingly, in multiple cases, we identified a subset of peptides that were phenotype-exclusive even though the protein was not. While we have not ruled out the possibility that signal interference or mass spectrometric limits of detection could explain these binary differences, we suspect that disease-specific contexts may be altering downstream proteolysis and presentation. Phenotype-specific proteolysis could be important in identifying conditionally distinct peptide products, which would further broaden the scope of disease-relevant targets, and showcases these perturbations have widespread impacts on the immune repertoire. Indeed, recent work has shown that drugs can induce significant changes in the immunopeptidome and enriched HLA-peptide complexes can be targeted with recombinant antibodies for more selective killing^25^.

In conclusion, the sHLA method can pull out HLA-associated peptides in an allele-restricted, phenotype-specific manner that matches immunopeptidomes obtained by more laborious methods. The growing interest in development of antibody or T-cell receptor-based therapeutics against specific disease-associated HLA-peptide complexes depends critically on the identification of allele-specific immunopeptidomics for target discovery and peptide triage. We believe this information of disease-associated peptides or neoantigens may accelerate personalized medicine and reshape how we assess the potential of certain antigens.

## METHODS

### Cloning

sHLA Fc-fusion was cloned into the lentiviral vector pLVX-TetOne-Puro. All constructs were sequence verified by Sanger sequencing.

### Cell lines and culturing

B721.221 cells were a generous gift from Dr. Lewis Lanier (UCSF). Capan-1 pancreatic cancer cells were from Wells lab frozen stocks. KP4 and MiaPaCa-2 pancreatic cells were generous gifts from Dr. Rushika Perera (UCSF). A549 lung cancer cells were a gift from Dr. Oren Rosenberg (UCSF). PC3 prostate cancer cells and MCF7 breast cancer cells were purchased from the UCSF Cell Culture Facility. B721.221, A549, PC3, and MCF7 cells were all grown in RPMI+10% Tetracycline-negative FBS+1% Pen/Strep. Capan-1, KP4, and MiaPaCa-2 cells were all grown in IMDM+10% Tetracycline-negative FBS+1% Pen/Strep. Cell lines transduced with the sHLA Fc-fusion were cultured in media with 2 μg/mL puromycin. sHLA Fc-fusion cell lines were cultured in respective media without FBS and with doxycycline for sample collection. All cells were grown at 37°C, 5% O_2_ unless otherwise stated.

To generate normoxic Capan-1, KP4, and MiaPaCa-2 cells, cells were grown for 3 days in IMDM+10% tetracycline-negative FBS+1% Pen/Strep at 37°C, 5% O_2_ before beginning doxycycline treatment. Hypoxic cells were grown in a hypoxic chamber in IMDM+10% Tetracycline-negative FBS+1% Pen/Strep at 37°C, 1% O_2_ prior to doxycycline treatment. Hypoxic cells were only removed from the chamber to replace media for the appropriate condition and the exchange was conducted as quickly as possible to avoid the onset of the normoxic phenotype.

A549, MCF7, and PC3 cells were seeded one day prior to treatment. Cells were incubated in media containing either 250 nM doxorubicin (Sigma-Aldrich) or the equivalent volume of DMSO for 24 hours. Growing samples were treated with doxycycline immediately after DMSO treatment. For the senescent samples, media was replaced and then subsequently replaced every other day for 8 days post-doxorubicin treatment before doxycycline treatment. Cells were seeded separately for western blot analysis and β-galactosidase activity staining. β-galactosidase activity staining was performed using a Senescence β-Galactosidase Staining Kit (Cell Signaling) following the manufacturer’s protocol.

### Lentivirus and cell line generation

HEK293T cells were cultured in DMEM+10% FBS+1% Pen/Strep. Cells were seeded 5×10^5^ per well of a 6-well plate a day prior to transfection. Plasmids at the designated concentrations (1.35 μg pCMV delta8.91, 0.165 μg pMD2-G, 1.5 μg sHLA plasmid) were added to OptiMEM media with 9 μL FuGENE HD Transfection Reagent (Promega) at a 3:1 FuGENE:DNA ratio, incubated for 30 minutes, and subsequently transfected into HEK293T cells. The supernatant was harvested and cleared by passing through a 0.45-μm filter 72 hours post transfection. Cleared supernatant was added to target cells (∼1 million cells) with 8 μg/mL polybrene and centrifuged at 1000 x g at 33°C for 3 hours. 24 hours post-transduction, media was replaced with appropriate fresh media. After an additional 24 hours, drug selection for stable cell lines was initiated by the addition of 2 μg/mL puromycin and expanded.

To expand successfully transduced B721.221 cells, live cells were isolated using SepMate™-50 (IVD) tubes and Lymphoprep™ (Stemcell Technologies). For isolation, cell cultures were centrifuged at 300 x g for 5 minutes and resuspended in 5 mL of cell culture media. 15mL Lymphoprep™ was added to each SepMate™-50 (IVD) tube, and the 5 mL cell suspension was subsequently added. Tubes were centrifuged at 400 x g for 10 minutes, and then supernatant was quickly decanted into 30 mL cell media and the SepMate™-50 (IVD) tube was discarded. The cell culture was spun at 300 x g for 5 minutes and supernatant was removed. Pellets were resuspended in cell media containing appropriate drug and expanded. A total of 2 isolations occurred for each cell line.

### Sample Preparation

Cell phenotypes were induced to produce the sHLA-Fc fusions as described. Cells were cultured in media with 2 μg/mL doxycycline for 24 hours, and then subsequently cultured in serum-free media with 2 μg/mL doxycycline for 28 hours prior to media collection. For each 50 mL of media sample, 100 μL of Pierce™ Protein A Magnetic beads (Thermo Scientific) were washed twice with PBS prior to use. Beads were added to media—which has previously been filtered with 0.45-μm filters—and rotated for 1 hour at 4°C. Samples were spun at 500 x g for 5 minutes and media removed. Beads were washed strenuously with 10 mM Tris pH 8.0 made with Optima™ LC/MS water (Thermo Scientific). After washing, protein/peptides were eluted by incubating beads with 10% acetic acid for 10 minutes at room temperature. Beads were washed twice with 10% acetic acid, and washes and elution were pooled together. Samples were dried in a Genevac prior to desalting.

Dried down samples were resuspended in 75 μL of 1% TFA and vortexed vigorously. Samples were centrifuged at 21,000 x g for 5 minutes at RT to remove any remaining precipitate. Sample was placed in a magnetic rack, after which the supernatant was removed gently from the tube and placed in a prepared Pierce C18 column as per manufacturer’s instruction. Shortly, each column was washed with 200 μL of 70% acetonitrile in water and spun down at 1500xg until dry. The columns were further washed with 200 μL of 50% acetonitrile in water and spun till dryness. Following the pre-wash steps, each column was further washed twice with 200 μL of 5% acetonitrile/0.5% TFA in water and spun till dryness. The sample was then loaded onto the column and spun till dryness. Each sample was reloaded onto the column to maximize peptide yield. Samples were then washed with 2x 200 μL of 5% acetonitrile/0.5% TFA in water, 200 μL of 5% acetonitrile/1% FA in water, and eluted in 2x 50 μL of 70% acetonitrile in water. Samples were dried to completion.

### Mass Spectrometry

Liquid chromatography and mass spectrometry was performed as previously described^26^. Briefly, each sample was brought up in 6.5 μL of 2% acetonitrile/0.1% formic acid in water, vortexed vigorously, and spun down at maximum speed to remove any precipitate. The sample was transferred and 6 μL of the peptide supernatant was separated using a nanoElute UHPLC system (Bruker) with a pre-packed 25 cm x 75 μm Aurora Series UHPLC column + CaptiveSpray insert (CSI) column (120 A pore size, IonOpticks, AUR2-25075C18A-CSI) and analyzed on a timsTOF Pro (Bruker) mass spectrometer. Peptides were separated using a linear gradient of 7-30% (Solvent A: 2% acetonitrile, 0.1% formic acid, solvent B: acetonitrile, 0.1% formic acid) over 60min at 400 nL/min. Data-dependent acquisition was performed with parallel accumulation-serial fragmentation (PASEF) and trapped ion mobility spectrometry (TIMS) enabled with 10 PASEF scans per top N acquisition cycles. The TIMS analyzer was operated at a fixed duty cycles close to 100% using equal accumulation and ramp times of 100 ms each. Singly charged precursors below 800m/z were excluded by their position in the m/z-ion mobility plane, and precursors that reached a target value of 20,000 arbitrary units were dynamically excluded for 0.4 min. The quadrupole isolation width was set to 2 m/z for m/z < 700 and 3 m/z for m/z > 700 and a mass scan range of 100-1800 m/z. TIMS elution voltages were calibrated linearly to obtain the reduced ion mobility coefficients (1/K0) using three Agilent ESI-L Tuning Mix ions (m/z 622, 922, and 1,222).

### Data Analysis

Briefly, for general database searching, peptides for each individual dataset were searched using PEAKS Online X version 1.5 against the entire Swiss-prot Human Proteome (Swiss-prot). Enzyme specificity was set to Unspecific. Peptide length was specified between 8-12 amino acids. No fixed modifications were set, while acetylation (N-term) and methionine oxidation were set as variable modifications. The precursor mass error tolerance was set to 20 PPM and the fragment mass error tolerance was set to 0.03 Da. Data were filtered at 1% for both protein and peptide FDR. All mass spectrometry database searching was based off of four biological replicates. Biological replicates underwent preparation, washing, and downstream LC-MS/MS preparation separately. ICE Logos were generated using the server https://iomics.ugent.be/icelogoserver/. Novel peptide ID’s were determined by cross-referencing the Immune Epitope Database (as of 10/10/21).

### Western Blot

sHLA Fc-fusion samples from B721.221 cells were generated and purified as described. Beads were washed three times with PBS and protein was eluted with 0.1 M acetic acid. For growing and senescent samples, cells were washed twice on plate with PBS prior to lysis. Lysis buffer contained 1x RIPA (EMD Millipore), 1% protease inhibitor cocktail (Sigma-Aldrich), and 1 mM EDTA. Cells were lysed for 20 minutes on ice prior to sonication (1 minute, 20% amp, 1 second on/off pulse). Cells were spun at 16000 x g at 4°C for 5 minutes, and lysate protein concentration was determined using a Pierce™ BCA Protein Assay (Thermo Scientific). For hypoxic and normoxic samples, cells were washed twice on plate with PBS prior to dissociated by addition of versene (PBS+0.05% EDTA). Cells were lysed using lysis buffer containing 1x RIPA (EMD Millipore), 1% protease inhibitor cocktail (Sigma-Aldrich), and 1 mM EDTA. Cells were lysed for 20 minutes on ice prior to sonication (1 minute, 20% amp, 1 second on/off pulse). Cells were spun at 16000 x g at 4°C for 5 minutes, and lysate protein concentration was determined using a Pierce™ BCA Protein Assay (Thermo Scientific). Samples were run on a Bolt 4-12% Bis-Tris gel (Invitrogen), and transferred to a PVDF membrane (Thermo Scientific) using an iBlot™ (Thermo Scientific). Membranes were blocked in Odyssey® Blocking Buffer (TBS) (LiCOR) prior to staining. Membranes were stained with primary anti-FLAG (Cell Signaling, 14793S), anti-human HLA-A (Thermo Scientific, PA5-29911), anti-human HLA-B (Proteintech, 17260), anti-human Glut1 (Abcam, ab115730), and anti-human α-tubulin (Sigma-Aldrich, T6199) antibodies in blocking buffer for 1 hour at room temperature or overnight at 4°C. Secondary staining was performed using goat anti-rabbit IRDye® 800CW and goat anti-rabbit IRDye® 680RD antibodies (LiCOR Biosciences) in blocking buffer for 1 hour at room temperature. Membranes were washed with three 5 minute washes of TBST between each staining step. Membranes were imaged using an Odyssey® CLx (LiCOR Biosciences).

## Supporting information

sHLA Peptide Datasets

## ACKNOWLEDGEMENTS

We thank the members of the Wells lab for their support and Forest White for input on the manuscript. N.J.R was supported by the National Science Foundation Graduate Research Fellowship Program. L.L.K was s supported by the NIH F31 Ruth L Kirschstein National Research Service Award (1F31CA247527). J.A.W. was supported by generous funding from the Chan Zuckerberg Biohub 561 Investigator Program, the Harry and Dianna Hind Professorship, NIH (R35GM122451), and NCI 562 (R01CA248323).

## ABBREVIATIONS

(sHLA): secreted human leukocyte antigen
(MHC): major histocompatibility complex
(HLA): human leukocyte antigen
(β2M): β2-microglobulin

